# Eribulin normalizes pancreatic cancer-associated fibroblasts by simulating TGFβ inhibition

**DOI:** 10.1101/2022.02.25.482033

**Authors:** Tiffany Luong, Edna Cukierman

## Abstract

Less than 11% of pancreatic cancer patients survive 5-years post-diagnosis. The unique biology of pancreatic cancer includes a significant expansion of its desmoplastic tumor microenvironment, wherein cancer-associated fibroblasts (CAFs) and their self-produced extracellular matrix are key components. CAF functions are both tumor-supportive and tumor-suppressive, while normal fibroblastic cells are solely tumor-suppressive. Knowing that CAF-eliminating drugs are ineffective and can accelerate cancer progression, therapies that “normalize” CAF function are highly pursued. Eribulin is a well-tolerated anti-microtubule drug used to treat a plethora of neoplasias, including advanced/metastatic cancers. Importantly, eribulin can inhibit epithelial to mesenchymal transition via a mechanism akin to blocking pathways induced by transforming growth factor-beta (TGFβ). Notably, canonical TGFβ signaling also plays a pivotal role in CAF activation, which is necessary for the development and maintenance of desmoplasia. Hence, we hypothesized that eribulin could modulate, and perhaps “normalize” CAF function. To test this premise, we used a well-established *in vivo*-mimetic fibroblastic cell-derived extracellular matrix (CDM) system and gauged the effects of eribulin on human pancreatic CAFs and cancer cells. The pathophysiologic cell-culturing system was also used to query eribulin effects on CDM-regulated cancer cell survival and invasive spread. Results demonstrated that intact CDMs modestly restricted eribulin from obstructing cancer cell growth. Nonetheless, eribulin-treated CAFs generated CDMs that limited cancer cell survival under nutritional stress, similar to reported tumor-suppressive CDMs generated by TGFβ-deficient CAFs. Data from this study support the central proposed premise and suggest that eribulin could be used as a CAF-normalizing drug.

## Introduction

Pancreatic ductal adenocarcinoma (PDAC) is a devastating disease with limited therapeutic options [1]. This disease is estimated to become the second leading cause of cancer-related deaths in the United States by 2026 [2]. Currently, less than 11% of patients diagnosed with PDAC are estimated to survive up to 5-years [3]. Thus, it is of utmost importance to identify novel effective therapies for treating patients diagnosed with PDAC.

One of the major features of PDAC is its abundant desmoplastic stroma, characterized by an expansion of activated resident fibroblasts and the deposition of an interstitial extracellular matrix (ECM) by these cells. Since desmoplasia can encompass over 70% of the tumor mass, this unique tumor microenvironment (TME) plays a major role in the manner drugs influence disease progression [4-6]. Although the TME is known to modulate PDAC onset and progression, ablating the tumor stroma can accelerate tumorigenesis [7-9]. Consequently, efforts to prevent eroding the TME, while modulating it to attain a tumor-suppressive stroma, are highly sought [6, 10-14]. A possible approach for achieving cancer-associated fibroblast (CAF) “normalization” would be to effectively limit signaling induced by transforming growth factor-beta (TGFβ). This is because ligands from the TGFβ family are known to drive CAF activation [10]. Further, high levels of phosphorylated Smad2/3 (pSmad2/3; Small Worm Phenotype and Mothers Against Decapentaplegic 2 and 3) in human CAFs, which are indicative of constitutively high canonical TGFβ activity in these cells, have been shown to inversely correlate with PDAC and renal cancer overall survival [15]. Unfortunately, directly targeting TGFβ has not been clinically efficacious [16]. Hence, drugs that simulate TGFβ inhibition, yet demonstrate high patient tolerability (e.g., imparting low toxicity), could serve as valuable alternative therapeutics.

Eribulin is an anti-microtubule agent that was approved in 2010 by the Food and Drug Administration to treat metastatic breast cancer [17, 18]. A mechanism often used by metastatic cells that renders cells with stem-like properties is the TGFβ-modulated epithelial-to-mesenchymal transition (EMT [19]). Of note, eribulin was reported to revert metastatic EMT through a mechanism that dampens p-Smad2/3 levels [20]. Further, drugs known to prevent EMT, as well as drugs that block or revert mesenchymal cell activation [21], constitute potential stroma-normalizing agents [10, 13-15, 22-26]. Hence, we hypothesized that treating CAFs with eribulin could simulate TGFβ-signaling blockage and render a “normalized” (e.g., tumor-suppressive) microenvironment.

To this end, our team previously developed an *in vivo*-like fibroblastic three-dimensional (3D) cell culturing system [27, 28], and has adapted it for the use of PDAC patient-harvested CAFs [15, 22, 23, 29]. This system can be used for assessing how drugs modulate CAF function, as well as how drug effects are altered by fibroblastic ECMs [28, 30, 31]. Hence, our CAF-generated <u>c</u>ell-<u>d</u>erived EC<u>M</u> system, known as “the CDM system,” was used in this study for conducting a series of tests that were designed to measure the effects of eribulin in modulating CAF phenotypes and functions. Further, since pancreas resident fibroblastic cells, initially presenting with normal-like tumor-suppressive features, undergo an ex-vivo culturing stress-induced activation [32], our team previously used the CRISPR/Cas9 system to engineer β5-integrin deficient PADC CAFs (β5^KO^) [15]. These β5^KO^ CAFs, serve as “normalized” CAF controls [22, 33], as these cells perpetuate a phenotype that simulates TGFβ inhibition in CAFs [15] and constituted an important model used in this study. We herein report that CAF-generated ECMs protect PDAC cells by modestly hindering the effects of eribulin upon limiting tumor cell growth. Importantly, we aim to report that eribulin also acts directly on CAFs and prompts these cells to generate an ECM that no longer effectively promotes PDAC cell invasive spread and has lost its ability to nurture PDAC cells under nutritional stress. Of note, the observed CAF-generated ECM functional shift, from tumor-supportive to tumor-suppressive, simulated reported results that were obtained with TGFβ deficient CAFs in our CDM system [15, 22, 33]. Hence, this study proposes that eribulin could be used as a stroma-normalizing agent.

## Materials and Methods

### Ethics statement informing on the previously collected tissue used to harvest the human fibroblastic cells utilized

All fibroblastic human cells used in this study, were harvested from surgical patient samples in accordance with guidelines and regulations under the journal policies. Fresh surgical samples were collected by the Biosample Repository Facility (BRF) at Fox Chase Cancer Center. Prior to surgery, patients interested in donating tissue for research are informed on how the tissue is collected, processed, and distributed, as well as the type of data and decoding process that is undertaken by the BRF personnel. If patients agree, they proceed to sign a written consent that was approved by the Institutional Review Board (IRB) protocol at Fox Chase. To assure protecting the patient identities and specifically for this study, solely the BRF-generated specimen number and organ of origin were conveyed to the research team.

### Cells and culturing conditions

Human PDAC CAF lines were generated from two independent PDAC surgical samples. Using the parental CAF line from one of these patients, a GFP-CRISPR^KO^ control was also used and it is referred to as CAF’ throughout the text. CAF’ served as a control for the β5-integrin deficient CAF line (β5^KO^), which was engineered from the same parental CAF line. Of note, β5^KO^ CAFs are known to sustain normal-like traits and functions. This is because these normalized CAFs cannot be activated in the absence of a mature TGFβ1 ligand [15]. A matching tumor-adjacent fibroblastic CAF line maintaining normal-like traits and functions, and harvested from a benign PDAC adjacent tissue, was used in the re-plating experiments designed to test the ability of CAF-generated ECMs to induce a normal-to-CAF activation overnight, as published [15, 23]. CAFs harvested by a second PDAC surgical sample are referred to as “CAF 2” throughout the text. Additionally, a human lung CAF line was used for added rigor. All fibroblastic cells were isolated from fresh surgical tissue collected at Fox Chase Cancer Center (Philadelphia, PA) using an enzymatic tissue digestion approach, and immortalized with hTERT as published [28]. Fibroblasts were authenticated and tested for mycoplasma as previously reported [15, 23]. Panc-1 human PDAC cells were purchased from the American Type Culture Collection (Manassas, VA) and used within 6 months of thawing. All cells were cultured in high-glucose Dulbecco’s Modified Eagle Medium (DMEM) supplemented with 10% Fetal Bovine Serum (FBS, Peak Serum, Wellington, CO), 2 mM L-glutamine, and 100 U/ml-µg/ml and penicillin-streptomycin (Corning, Manassas, VA) at 37°C with 5% CO_2_ unless otherwise specified.

### Chemicals or Drugs

Eribulin-mesylate was provided as a lyophilized powder by Eisai Inc. (Woodcliff Lake, NJ). Lyophilized powder was reconstituted in HPLC grade DMSO to make 10 mM stock solutions. Paclitaxel was purchased from Sigma-Aldrich (St. Louis, MO) and reconstituted in DMSO to make 1.17 mM stock solutions. Stock solutions were aliquoted for single-use thawing, stored at -80°C, and diluted with cell culture medium to achieve the desired experimental concentrations. SB431542 hydrate, a selective inhibitor of TGFβ Type 1 receptor kinases (TGFR1i) was purchased from Sigma-Aldrich (St. Louis, MO) and used at a concentration of 10 µM in all experiments.

### Viability Assay (2D)

Panc-1 or CAFs were seeded at a density of 2 × 10^3^ cells per well in a 96-well plate, overnight. The next day, cells were treated with either eribulin, paclitaxel, TGFR1i, or DMSO daily for 5 days at the indicated concentrations. Alamar blue (Thermo Fisher Scientific, Waltham, MA) was added on the 6^th^ day and incubated at 37°C, 5% CO_2_ for 4 hours. Cell viability was measured using Tecan Spark™ 10M microplate reader (Tecan, Switzerland) at an absorbance of 570 nm and a reference wavelength of 600 nm according to the manufacturer’s protocol.

### Fibroblastic cell-derived ECM (CDM) system

Fibroblastic cell-derived ECMs (known as CDMs) and decellularized CDMs were generated as previously published [28, 34]. The CDM system was used to measure ECM production in the presence of eribulin or drug controls (e.g., paclitaxel, TGFR1i, or DMSO vehicle) at the concentrations stated in the text. Briefly, confluent fibroblastic cultures were supplemented with 50 µg/mL of ascorbic acid daily for a total of 5 days [28, 34]. The quality of ECM production was confirmed via indirect immunofluorescence (see method below). The thickness of ECM, indicative of qualified matrix production, was measured to reach a minimum of 7 µm; ECMs thinner than this were not considered of high quality and were hence excluded from all analyses. For re-plating and 3D survival assays (see method below), CDMs were decellularized by treating cultures with PBS containing 0.5% Triton X-100 and adding 20 mM NH_4_OH (Sigma-Aldrich, St. Louis, MO) at the end of ECM production. The resulting decellularized CDMs were treated with DNase I and washed with PBS before cellular re-inoculation, as published [28]. Eventually, tumor-adjacent fibroblasts or PDAC cells were re-plated.

### Normal-to-CAF functional transition: Re-plating Assay

Decellularized drug or vehicle-treated CAF-generated CDMs were used to re-plate tumor-adjacent fibroblastic cells, obtained from a PDAC adjacent tissue, matching one of the PDAC CAFs [15]. 7 × 10^3^ fibroblastic cells, from the above described adjacent tissue, vetted first to sustain normal-like traits [28, 34], were seeded onto the assorted CAF-generated CDMs and cultured overnight. Levels of αSMA co-localization with stressed fibers (F-actin, phalloidin staining) were measured, as these are indicative of an effective ECM prompted normal-to-CAF fibroblastic cell transition [15, 27]. Cultures were stained with αSMA at 1:300 (100 µg/ml) and phalloidin 1:40 (3.3 µM) via indirect immunofluorescence (see method below). Images were acquired with a plan Apo λ 60x oil immersion objective using a Nikon A1S confocal system. Monochromatic image stacks were reconstituted in ImageJ (NIH, Bethesda, MD) and the percentage levels of co-localization of αSMA to stressed fibers (F-actin, phalloidin stain) was assessed using the MetaMorph 7.8.0.0 software.

### Survival Assay (3D)

Survival assay was performed similarly to what was published by Francescone, R., et.al [23]. Briefly, 1.5 × 10^3^ red fluorescent protein (RFP)-expressing Panc-1 cells were seeded onto assorted decellularized CDMs under nutrient-depleted conditions (*i*.*e*., serum-free, L-glutamine-free media; unless otherwise stated), in the presence or absence of eribulin and compared with assorted drug and vehicle controls. The numbers of cells were measured/acquired every 24 hours, up to 120 hours, using a Nikon A1 confocal system equipped with a 10x (plan Apo) non-oil objective. Cell numbers were gauged as percentage red-cell area coverages per image with a minimum of 5 images acquired per sample.

### Cancer cell Invasion Spread Assay

ECM invasion spread assays were conducted as previously published [35]. Briefly, 3 × 10^3^ Panc-1 cells per 30 µl were pre-labeled with Hoechst 33342 (Thermo Fisher Scientific; Waltham, MA) for 1 hour prior to trypsinization and resuspended in tumor sphere-forming media (Irvine Scientific; Santa Ana, CA) supplemented with 2 U/ml of heparin and 0.5 µg/ml hydrocortisone. The individual 30 µl sample drops were carefully placed on a sterile 100 mm culture dish lid overnight with drops hanging upside down allowing spheroid formation. 5 ml of sterile PBS was added to the bottom of the dish to prevent hanging drops from drying out. Following 24 hours, spheroids were carefully collected and deposited onto assorted decellularized CAF-generated CDMs, incubated for a few minutes to allow spheroids to adhere, then media was carefully added to the culture. Spheroids were incubated for 2 hours to fully adhere, and this time was considered baseline (*i*.*e*., time 0). Images were acquired at baseline and again at 24 hours for each spheroid using 4x magnification. For each condition, the spread of the spheroids was measured and reported as mean fold invasive spread (mean radius) per sphere. A minimum of 3 spheroids were recorded per experimental condition and experiments were repeated a minimum of two times per condition.

### Indirect Immunofluorescence

Samples were labeled by indirect immunofluorescence as published [27, 28]. Briefly, experimental samples were generated/cultured onto 12 mm coverslips (Carolina Biological, Burlington NC). These were fixed and permeabilized in PBS supplemented with 4% paraformaldehyde (Electron Microscopy Sciences, Hatfield, PA), 5% sucrose (w/v), and 0.5% Triton X-100 for 3 minutes, followed by additional 20 minutes fixing with PBS containing 4% paraformaldehyde and 5% sucrose (w/v). Samples were blocked for a minimum of 1 hour with Odyssey Blocking Buffer-PBS (LI-COR Biosciences, Lincoln, NE). The following primary antibodies were incubated for 90 minutes at room temperature using the antibodies and concentrations listed below.

**Table 1.** List of indirect immunofluorescence primary antibodies.

| **1° Antibody/reagent** | **Host** | **Concentration** | **Source** | **Catalog#** |
| --- | --- | --- | --- | --- |
| αSMA | Mouse | 100 μg/ml | Sigma-Aldrich | A2547 |
| Fibronectin | Rabbit | 3 μg/μl | Sigma-Aldrich | F3648 |
| anti-pFAK Y<sup>397</sup> | Rabbit | 2.5 μg/μl | Invitrogen | 44-624G |
| anti-human active α5β1 integrin | Mouse | 45 μg/ml | SNAKA51, a gift from M Humphries [31] | N/A |
| DAPI | N/A | 0.25 μg/μl | Invitrogen | D1306 |

Samples were washed with PBS-Tween20 (0.05%) three times for 5 minutes each. Secondary donkey F(ab’) fragments [1:100] cross-linked to assorted fluorophores (Jackson ImmunoResearch Laboratories Inc., West Grove, PA), were incubated for 60 minutes at room temperature. Samples were washed as above and mounted using an anti-fade mounting reagent (Corning imaging recipe). Images were acquired with plan Apo λ 60x oil immersion objective using a Nikon A1S confocal system as above. Acquisition settings were kept identical for each primary antibody/fluorophore-liked F(ab’) secondary fragment. Sequential monochromatic images, corresponding to 0.5 µM thick slices, were reconstituted in ImageJ and processed for ECM or activated CAF biomarker evaluations as needed. A minimum of 5 images were acquired per sample. A minimum of 2 samples were prepared for each experiment, while experiments were repeated at least twice for each condition.

### Western Blot

CDM-system generated cell cultures were lysed in RIPA buffer containing the following solutions: 150 mM sodium chloride, 1.0% Triton X-100, 0.5% sodium deoxycholate, 0.1% SDS (sodium dodecyl sulphate), and 50 mM Tris, pH 8.0. Protease and phosphatase inhibitors (Thermo Fisher Scientific, Waltham, MA) were added fresh before use. Cell lysates were resolved by SDS-PAGE (Sodium Dodecyl Sulfate PolyAcrylamide Gel Electrophoresis) using 4-20% precast gel (Biorad, Hercules, CA). Proteins were transferred by semi-dry transfer onto a PVDF (polyvinylidene fluoride) membrane. Membranes were blocked using 5% non-fat milk (Biorad, Hercules, CA) in TBST (0.01% Tween20) for a minimum of 1 hour at room temperature. The following primary antibodies were incubated overnight at 4°C as indicated in the table below.

**Table 2.** List of western blot primary and secondary antibodies.

| 1° Antibody | Host | Dilution | Source | Catalog# |
| --- | --- | --- | --- | --- |
| anti-palladin | Rabbit | 0.0002 µg/µl | Protein tech,<br>Chicago, IL | 10853-1-AP |
| anti-phospho Smad2<br>[S465/467]/Smad3<br>[S423/425] | Rabbit | 0.2 µg/ml | Cell Signaling,<br>Danvers, MA | 8828s |
| Anti-mouse IgG, HRP-linked | Mouse | 1:5000 | Cell Signaling,<br>Danvers, MA | 7076S |
| Anti-Rabbit IgG, HRP-linked | Rabbit | 1:5000 | Cell Signaling,<br>Danvers, MA | 7074S |

Following overnight incubations, membranes were washed 3 times with TBST for 15 minutes each. HRP-linked secondary antibodies were added and incubated for 1 hour at room temperature. Membranes were again washed 3 times with TBST for 15 minutes each. Lastly, membranes were incubated with Immobilin Chemiluminescent HRP Substrate (Millipore, Burlington, MA) for 5 minutes before being developed onto films.

### Statistical Analysis

For statistics in all experiments, we used the one-way ANOVA test to compare between conditions; all asterisks indicate comparisons of specified conditions vs DMSO treatment (* p < 0.05, ** p < 0.01, *** p < 0.001, **** p < 0.0001) unless otherwise indicated in the figure legend. For comparison of fold growth rates in Figure 1C, a nonparametric t-test was used to compare between each pair data set at 120 hours. Note that for all (2D and 3D) culture conditions, effects of DMSO (e.g., vehicle) were used to normalize results and were compared to the behavior of untreated cells, finding no significant differences, unless otherwise noted.

**Figure 1.**
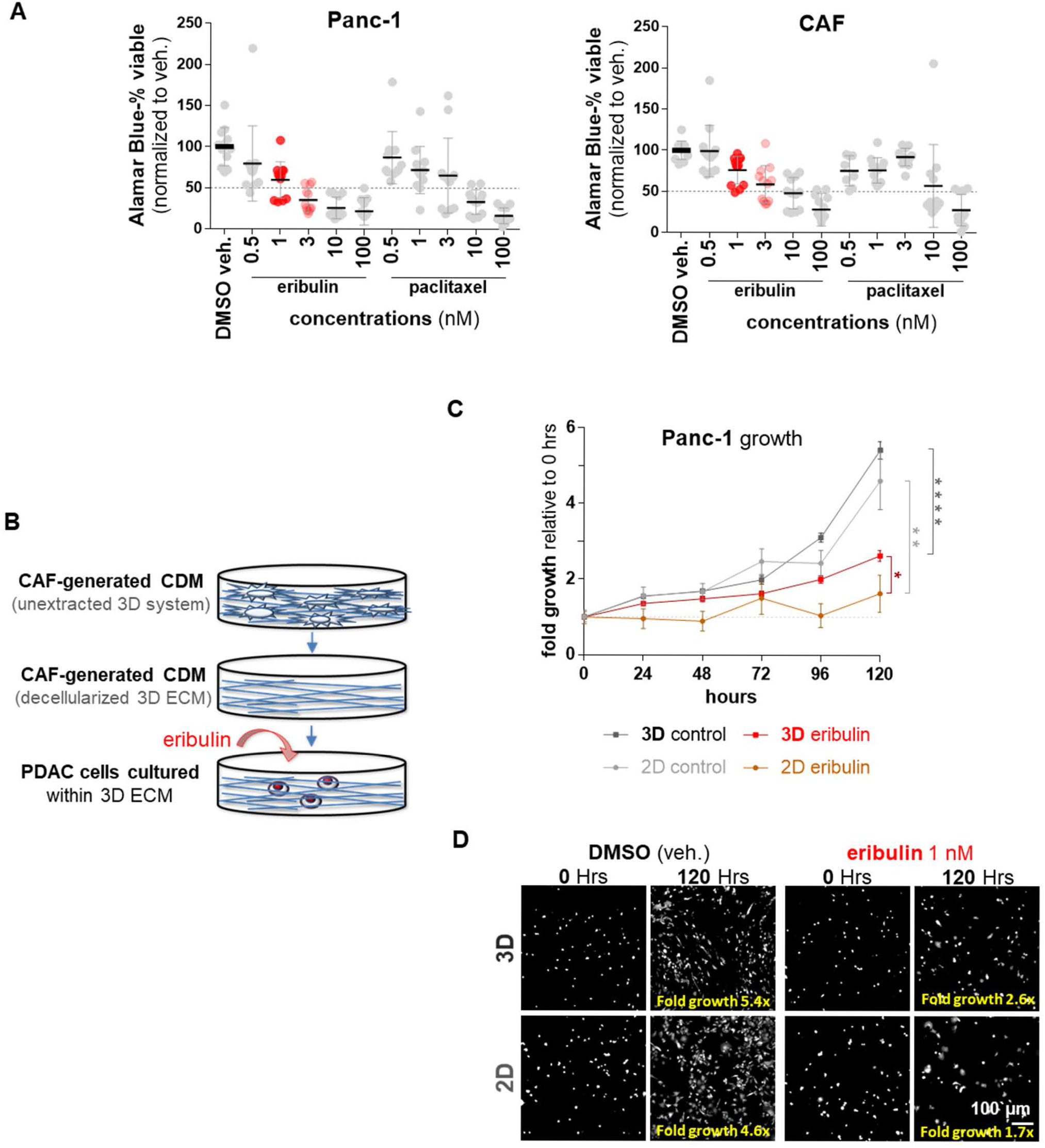
CAF-generated ECM protects PDAC cells from eribulin-induced growth inhibition. (A) Growth effects imparted by eribulin vs paclitaxel on Panc-1 vs. CAFs, measured via Alamar Blue assay, using classical 2D culturing conditions. Graphs show the percentage of viable cells, compared to DMSO vehicle control. Drugs were used at the stated concentrations for 5 days. Experiments were repeated 4 times using a minimum of 4 replicates per experimental condition. (B) Illustration depicting experimental conditions whereby RFP-expressing Panc-1 cells cultured within decellularized CAF-generated CDMs (blue) were treated daily with eribulin (red). (C) Graph indicating fold cell-growth values normalized to each condition at day 0 and measured every 24 hours for 120 hours. Values shown represent means and standard errors, while asterisks are indicative of a nonparametric t-test, used to compare between data sets at 120 hours; * P < 0.05; ** p < 0.01; *** P <0.001; **** p < 0.000. (D) Representative images of Panc-1 cell area coverages at baseline (time 0) and 120 hours, using 2D and 3D conditions showing DMSO vehicle and eribulin treatments. Fold area coverage increases are listed in yellow as fold growth. Scale bar corresponds to 100 µm (white).

## Results

### CAF-generated CDM protects PDAC cells from eribulin induced growth inhibition

Pre-clinical and clinical studies have demonstrated that CAF elimination could accelerate PDAC tumorigenesis [7-9]. Hence, in this study, it was initially important to assure that the eribulin needed half-maximal inhibitory concentration (IC_50_), intended to eliminate PDAC cells, was not significantly harmful to human PDAC CAFs. To test this, CAFs harvested from surgical samples [34] and the well-known PDAC cell line Panc-1 [36], were independently cultured, using classic two-dimensional (2D) conditions, in the presence of increased concentrations of eribulin. Results were compared to results obtained with paclitaxel, as this drug constitutes another anti-microtubule drug often used in the clinic, while DMSO served as vehicle control. Figure 1A shows that 5-day treatment with 1 nM and 3 nM eribulin rendered a 40% and 65% decline (mean 60% ± 22% and 35% ± 15%, respectively) in PDAC cell viability compared to vehicle control (DMSO mean 100% ± 23%). Interestingly, CAFs failed to reach 50% cell growth inhibition in the presence of eribulin at the same concentrations (76% ± 17% for 1 nM and 59% ± 23% for 3 nM) (Figure 1A).

It is important to note that the *in vivo*-mimetic three-dimensional (3D) CDM system is suitable for measuring ECM modulation of human cancer cell growth in response to chemotherapeutic drugs [30]. Therefore, we next questioned whether the rate of PDAC growth inhibition induced by eribulin could be altered in the presence of decellularized CAF-generated CDMs. For this, PDAC cell growth was gauged under classic 2D conditions, or within decellularized CAF-generated 3D CDMs (Figure 1B). Cells were treated daily with 1 nM eribulin or equal volumes of DMSO vehicle (*i*.*e*., control). PDAC cell growth values were calculated as area coverages, compared to baseline (e.g., day 0), every 24 hours for a total of 120 hours (e.g., 5 days). Results showed that PDAC cells grow at similar rates under both 2D and 3D conditions; area coverages measured in time presented with fit value linear regression slopes of about 0.03 for both 2D and 3D DMSO-treated cells (2.6 ×10^−2^ ± 6.0 × 10^−3^ and 3.2 × 10^−2^ ± 7.9 10^−3^, respectively). As expected, the slope value obtained for PDAC cells treated with eribulin under classic 2D conditions was the lowest (4.8 × 10^−3^ ± 2.5× 10^−3^). Indicating that the 2D condition resulted in the highest inhibition of PDAC cell growth, in response to eribulin. Notably, the decellularized CAF-generated 3D CDMs system (e.g., 3D substrate) provided a level of protection from eribulin with a growth slope of about 0.02 (± 1.7 × 10^−3^), which is 2.5-fold higher than its 2D counterpart (Figure 1C). Further, comparisons of initial PDAC cell area coverage at baseline vs. 120 hours, indicated that PDAC cells achieved ∼5-fold growth increases when treated with DMSO. Yet, area coverage of eribulin-treated PDAC cells cultured under 2D conditions failed to double after 5 days (1.7 fold), while the same cells cultured within the 3D CDMs increased the area coverage by almost 3-fold (Figures 1C and 1D). Altogether, these data indicate that decellularized ECMs, obtained using our CAF-generated CDM system, protect cancer cells from eribulin-imparted growth inhibition.

### Eribulin-treated CAFs simulate the traits of TGFβ-deficient CAFs

It is well known that fibroblastic cell traits and functions, *in vivo* and *in vitro*, are modulated and sustained by the self-generated interstitial ECM [37]. Hence, the above-listed results prompted to question whether the ECM-imparted protection against eribulin could be overcome using this drug to treat CAFs during ECM production (Figure 2A). We previously reported that blocking the TGFβ receptor, TGFR1, modulates CAF-generated CDM production. This is because TGFR1 inhibition renders a disorganized (e.g., isotropic) ECM with “normalized” features, which significantly diverges from the highly aligned (e.g., anisotropic) matrix fibers produced by intact tumor-supportive CAFs [15, 27]. Of note, it has been reported that in addition to acting as a microtubule dynamics-blocker [38], eribulin limits levels of pSmad2/3, which are indicative of obstructing canonical TGFβ signaling [20, 39]. The eribulin-modulated ECM features were hence compared to results attained with paclitaxel and with the small molecule SB431542, a well-known TGFR1 inhibitor (TGFR1i). In addition, the reported αvβ5-integrin deficient (β5^KO^) CAFs, known to perpetuate normalized CAF features (and functions) akin to TGFβ inhibition [15], served as a positive control indicative of “normalized” CAF-generated CDMs. To assure a high quality of ECM production via the CDM system, controls included measuring cell density and ECM thickness. Respectively, these tests assure that the 5-day lasting treatments do not significantly alter CAF numbers, or the ability of CAFs to uphold fibrillogenesis (Supplemental Figure 1A). Note that solely sufficiently thick ECM samples were used to assess fiber alignment (Figures 2B and Supplemental 1B). To this end, the “normalized” β5^KO^ CAF control exhibited both an expected modestly-low cell density [27], and a productive fibrillogenesis [15] (Supplemental Figure 1A). Similarly, results revealed that eribulin does not compromise ECM fibrillogenesis (Supplemental Figure 1A). CAFs treated with 3 nM paclitaxel attained poor fibrillogenesis, which invalidated the further assessment these ECMs (Figures 2B and Supplemental 1B). Regarding ECM alignment, results showed that eribulin significantly reduced the parallel fiber organization of CAF-generated CDMs, suggesting an ECM “normalization” (Figures 2B and Supplemental 1B). Further, these isotropic fiber disorganization levels were comparable to the levels obtained with TGFR1i, as well as in ECM fibers produced by the β5^KO^ CAFs (Figures 2B and Supplemental 1B). For rigor purposes, CAF’, the CRISPR/Cas9 control of the β5^KO^ CAFs, as well as an additional PDAC patient-harvested CAF line (CAF 2) and a lung cancer CAF line were also tested, and all rendered similar eribulin-modulated changes in CAF-generated CDM fiber alignment (Supplemental Figure 1B). Together these results suggest that eribulin effectively phenocopies the effects attained with TGFβ deficient CAFs (e.g., using TGFR1i treatment or in β5^KO^ CAFs) during ECM fibrillogenesis. Alternatively, treatment with paclitaxel either fails to alter ECM alignment, or if used at higher concentrations (e.g., 3 nM) compromises CAF fibrillogenesis.

**Figure 2.**
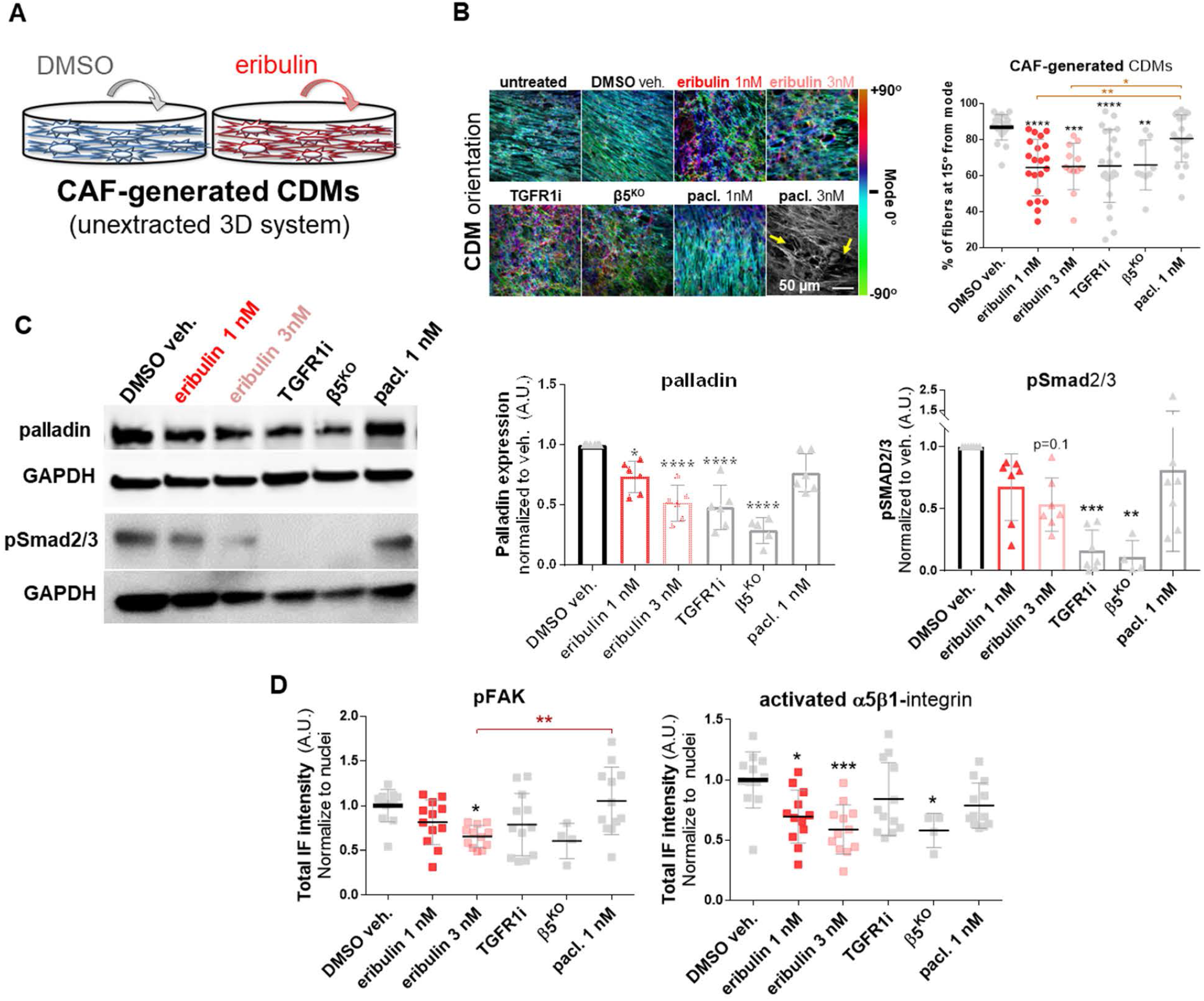
Eribulin normalizes CAF traits in the CDM system. (A) Illustration depicting the experimental conditions for CAF-generated CDMs that were treated with DMSO (blue) or eribulin (red) during the 5 days of matrix production. (B) Representative images of CAF-generated CDMs that resulted from the assorted experimental conditions used during the 5-day matrix production. Images show fibronectin fibers detected by indirect immunofluorescence and digitally pseudo-colored to depict the orientation angles of ECM fibers. The colored bar on the right represents the angle distributions and was normalized to show mode angle as 0° (cyan). The scale bar in the bottom right image corresponds to 50 μm. Note that CAFs treated with paclitaxel at 3 nM were not quantified due to the lack of fibrillogenesis as indicated by the yellow arrows pointing to ECM gaps and by the low cellularity and ECM thickness, measured below 7 µm, as shown in accompanying Supplemental Figure 1A. The graph indicates the levels of fiber alignment, quantified as the percentage of fibers oriented within 15° from the mode angle. (C) Representative immunoblots of lysates collected at the end of CAF-generated CDM production showing representative bands for palladin and phospho-Smad2/3 (pSmad2/3) using GAPDH as an intracellular protein loading control. Accompanying graphs depict levels of palladin and pSMAD2/3 gauged from immunoblots and normalized to levels attained under DMSO treated conditions. (D) The total intensity of phospho-FAK (pFAK) and Activated α5β1-integrin, measured using the MetaMorph software 7.8.0.0 measuring total intensities, with the same inclusive intensity thresholds set for all conditions and normalized to cell numbers (nuclei counts). All experiments were repeated 4 times while a minimum of 6 images were taken per sample. For statistical significance, one-way ANOVA was used to compare to DMSO vehicle. * P < 0.05; ** p < 0.01; *** P <0.001; **** p < 0.0001.

In addition to the effects of eribulin on ECM alignment, we questioned whether this drug could hinder known intracellular CAF biomarkers indicative of tumor-supportive CAF function [40]. For this, we quantified the levels of expression of the actin-bundling protein palladin, as well as constitutive pSmad2/3 activity. The two above-listed biomarkers are upregulated in tumor-supportive PDAC CAFs in response to TGFβ, and their intracellular fibroblastic levels inversely correlate with PDAC patient overall survival [15, 22, 41]. Similarly to using TGFR1i or β5^KO^ CAFs, increased doses of eribulin (e.g., 1 nM and 3 nM) triggered 30% and 50% decrease in palladin expression (0.73 ± 0.13 and 0.51 ± 0.14) respectively, with analogous tendencies for pSmad2/3 (0.67 ± 0.27 and 0.53 ± 0.21), compared to vehicle control (Figures 2C and Supplemental 1C). We also queried the effects of eribulin in the levels of focal adhesion kinase tyrosine-397 phosphorylation (pFAK), as well as intensities of the active conformation of the fibronectin receptor α5β1-integrin, as these two markers are too indicative of tumor-supportive CAF function *in vitro* (e.g., in the 3D CDM system), and have been associated with worse patient outcomes *in vivo* [15, 40]. Although levels of pFAK solely showed a modest tendency of 20% reduction in response to 1 nM eribulin (81% ± 25%), a significant decrease of 34% (66% ± 12%) in the activity of this kinase was evident with 3 nM eribulin (Figure 2D). Similarly, constitutive levels of active α5β1-integrin decreased by 30% and 40% in CAF-generating CDMs treated with 1 nM and 3 nM eribulin, respectively (Figure 2D). Note that all tendencies were similar to results obtained with TGFR1i and in β5^KO^ CAFs, yet paclitaxel did not affect the gauged pFAK or active α5β1-integrin levels. Altogether, these data suggest that eribulin triggers a phenotypic transition from tumor-supportive to tumor-suppressive, which is like known TGFβ inhibition and thus renders CAFs with features akin to “normalized” β5^KO^ CAFs [15, 40].

### Eribulin treated CAFs generate CDMs deficient in prompting a normal-to-CAF transition

Normal fibroblasts respond to CAF-generated CDMs by undergoing a phenotypic switch, analogous to CAF activation [27]. Since CDMs produced by eribulin-treated CAFs showed “normalized” features, we questioned whether these matrices have lost the functional ability to prompt a normal-to-CAF transition. For this, we used fibroblasts harvested from benign tissue adjacent to matching PDAC and seeded them onto decellularized CDMs (Figure 3A). Notably, decellularized CDMs obtained from eribulin-treated CAFs were no longer able to induce normal-to-CAF activation (Figure 3B), as denoted by the amount of αSMA localized to stress fibers [15, 27]. Although some basal αSMA levels were expressed by tumor-adjacent fibroblastic cells, the eribulin-treated CAF-generated decellularized CDMs solely induced about 2% of this protein to localize at stress fibers (median = 0.016 ± 0.28), compared to levels prompted by CDMs generated from CAFs treated with DMSO. These profiles were similar to the ones reported for CDMs obtained from TGFR1i treated or β5^KO^ CAFs, which too failed to induce CAF activation [15] (Figure 3B). Paclitaxel-treated CAFs, however, generated CDMs that were able to guide an αSMA stress fiber localization 10-fold higher than the one observed with eribulin (median = 0.2 ± 0.36) (Figure 3B). Altogether, results suggested that eribulin treatment of CAFs during CDM production renders matrices deficient in inducing a classic normal-to-CAF activation.

**Figure 3.**
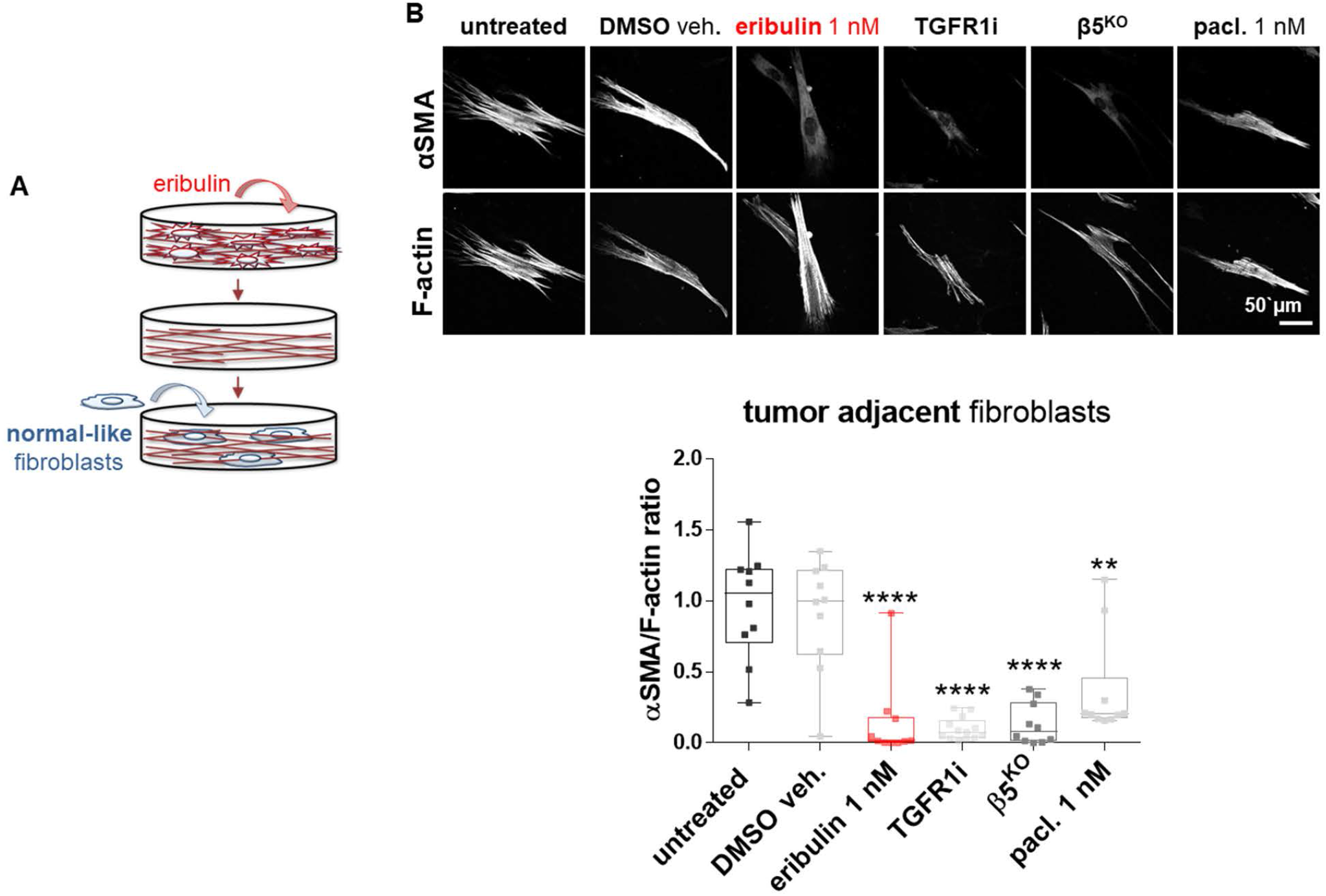
CAFs treated with eribulin generate ECMs that fail to induce a normal-to-CAF transition. (A) Illustration showing naïve fibroblasts re-plated into decellularized CDMs obtained from eribulin-treated CAFs. (B) Representative confocal immunofluorescent images showing monochromatic reconstitutions of αSMA and stress fibers (F-actin). Accompanying box and whiskers graph representing ratios of αSMA localization at F-actin positive pixels (error bars = maximum and minimum values while all median measured values are shown as individual squares). For statistical significance, ordinary one-way ANOVA was used comparing all data to the DMSO veh. control. * P < 0.05; ** p < 0.01; *** P <0.001; **** p < 0.0001.

### Eribulin treated CAFs generate CDMs deficient in promoting PDAC invasion and survival

We previously reported that similar to CDMs produced by normal fibroblasts, TGFβ-deficient CAFs (e.g., TGFR1i-treated and β5^KO^ CAFs) render CDMs that are compromised in promoting PDAC cell invasive spread and in sustaining PDAC cell survival under nutrient-deprived conditions [22, 23, 35]. Hence, we next questioned whether eribulin-treated CAFs would generate CDMs that too fail to sustain these pro-PDAC cell functions (Figure 4A). Using decellularized CDMs as before, results showed that when PDAC spheroids were cultured within CDMs produced by eribulin-treated CAFs for 24 hours, the invasive spreading capacity of the cancer cells was reduced by 20%, compared to vehicle-treated CAF-generated CDM controls (Figure 4B).

**Figure 4.**
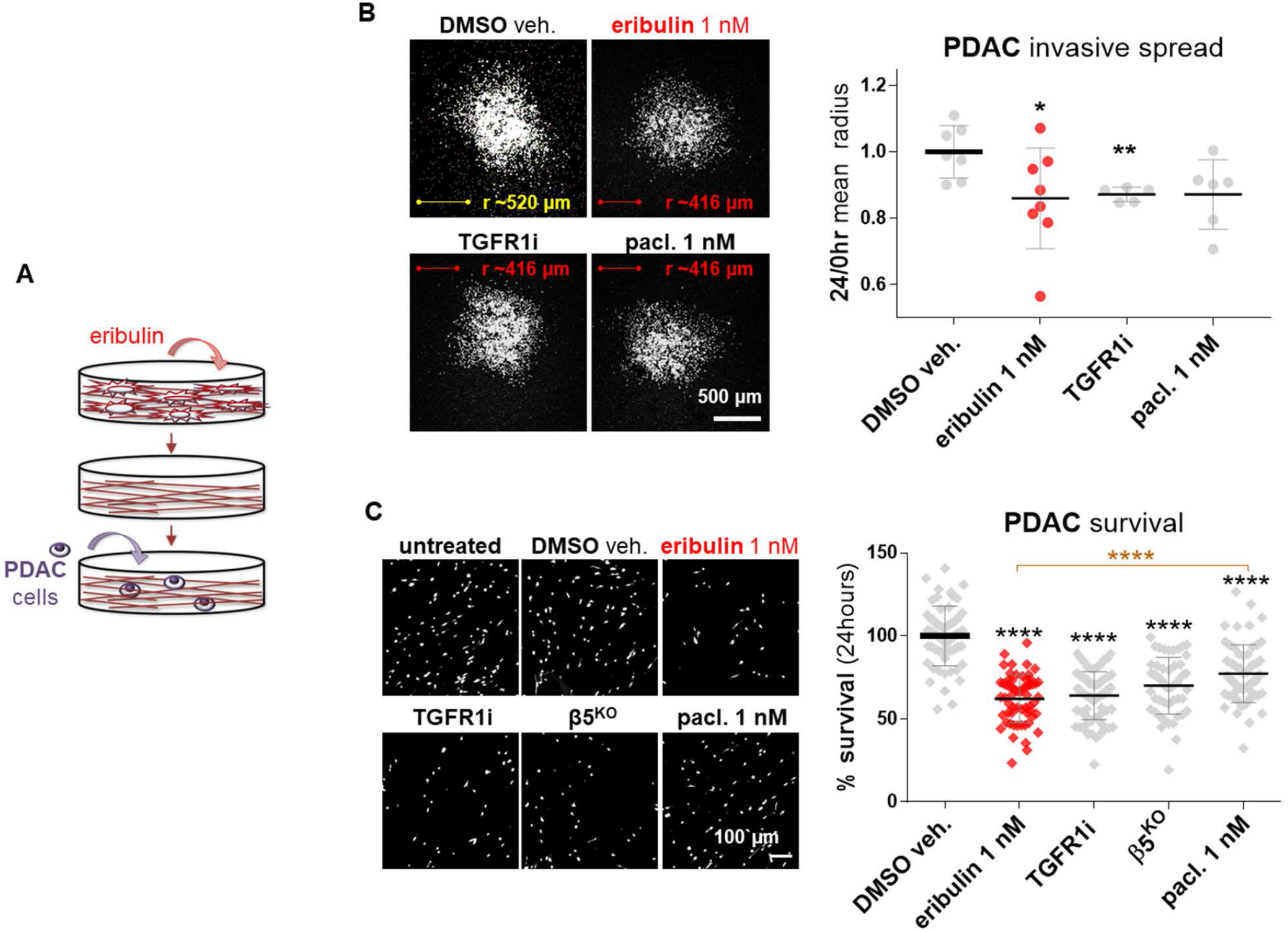
Eribulin-treated CAFs generate tumor-suppressive ECMs. (A) Illustration showing cancer cells re-plated onto decellularized CDMs obtained from eribulin-treated CAFs. (B) Representative 4x microscopy images of RFP-expressing Panc-1 spheroids that were allowed to spread for 24 hours (using nutrient-avid conditions) onto CDMs generated by CAFs treated with DMSO, eribulin, TGFR1i, or paclitaxel. Scale bar corresponds to 500 µm (white), while average spheroid spread radiuses (r) are shown in yellow for control and red for experimental conditions. The corresponding graph, on the right, shows 24hr/0hr ratios of measured mean-radiuses. For statistical significance, the Mann-Whitney test was used to compare each condition to the DMSO veh. control. * P < 0.05; ** p < 0.01; *** P <0.001; **** p < 0.0001. (C) RFP-expressing Panc-1 cells were cultured into the assorted listed decellularized CDMs, as in B, only this time using nutrient-deprived conditions. Viable cells were gauged following 24, 48, and 72 hours and representative 24-hour images are shown, while 48 and 72 hours are in Supplemental Figure 2. Scale bar corresponds to 100 µm. The graph indicates the percentage of survival, represented by RFP-expressing Panc-1 cells. Area coverage obtained data were normalized to DMSO Veh., which average was set as 100% survival. For statistical significance, a one-way ANOVA (Turkey’s multiple comparisons) test was used to compare all conditions. Black asterisks denote comparisons to DMSO, while the eribulin vs paclitaxel comparison is noted in orange. * P < 0.05; ** p < 0.01; *** P <0.001; **** p < 0.0001.

Importantly, PDAC cell survival assessed under nutrient-deprived conditions at 24, 48, and 72 hours, was significantly compromised (Figures 4C and Supplemental 2). After only 24 hours of culturing cells within CDMs generated by eribulin-treated CAFs, PDAC survival was decreased by 38% (± 14.6%) compared to the vehicle-treated control. Of note, these survival levels were also significantly limited by CDMs generated from TGFR1i-treated (36% ± 14.4%) and β5^KO^ (30% ± 17.1%) CAFs. Paclitaxel-treated CAFs generated CDMs that were solely able to modestly limit PDAC cell survival (23% ± 17.4%; Figure 4C). These results demonstrate that CAFs treated with eribulin produced CDMs incapable of providing a nutritional benefit to starved PDAC cells. Taken together, these data suggest that eribulin treatment effectively hinders CAFs’ ability to produce pro-tumoral CDMs, similar to matrices derived from normal-like fibroblasts [23].

### Eribulin fails to revert the survival benefit provided by intact CAF-generated CDMs

Results above (Figure 4C and Supplemental 2) demonstrated that CAFs treated with eribulin produced CDMs that limited PDAC cell survival under nutrient-deprived conditions. Hence, we were interested to test whether eribulin could also revert the benefit imparted by intact CAF-generated CDMs in sustaining PDAC cell survival under the same nutrient-poor conditions. For this, PDAC cells were cultured for 24 hours within decellularized but otherwise intact CAF-generated CDMs under nutrient-deprived conditions and treated with either eribulin, TGFR1i, paclitaxel, or a DMSO (Figure 5A). Results were compared to survival levels obtained using known tumor-suppressive CDMs (e.g., generated by TGFR1i-treated or β5^KO^ CAFs, as the ones used in Figure 4C). Of note, eribulin treatment failed to revert the protective effect imparted by decellularized CDMs (Figure 5B). Likewise, neither TGFR1i nor paclitaxel treatments hindered this CDM pro-survival benefit. These results suggest that eribulin effects on PDAC survival (e.g., Figures 4C and Supplemental 2), are the result of the eribulin-imparted effects on CAFs, as opposed to a direct effect upon PDAC cells. Altogether these observations are akin to reported TGFβ-deficient CAFs, whereby TGFβ signaling was shown to be uniquely necessary to produce a tumor-supportive ECM [15, 22, 33, 40].

**Figure 5.**
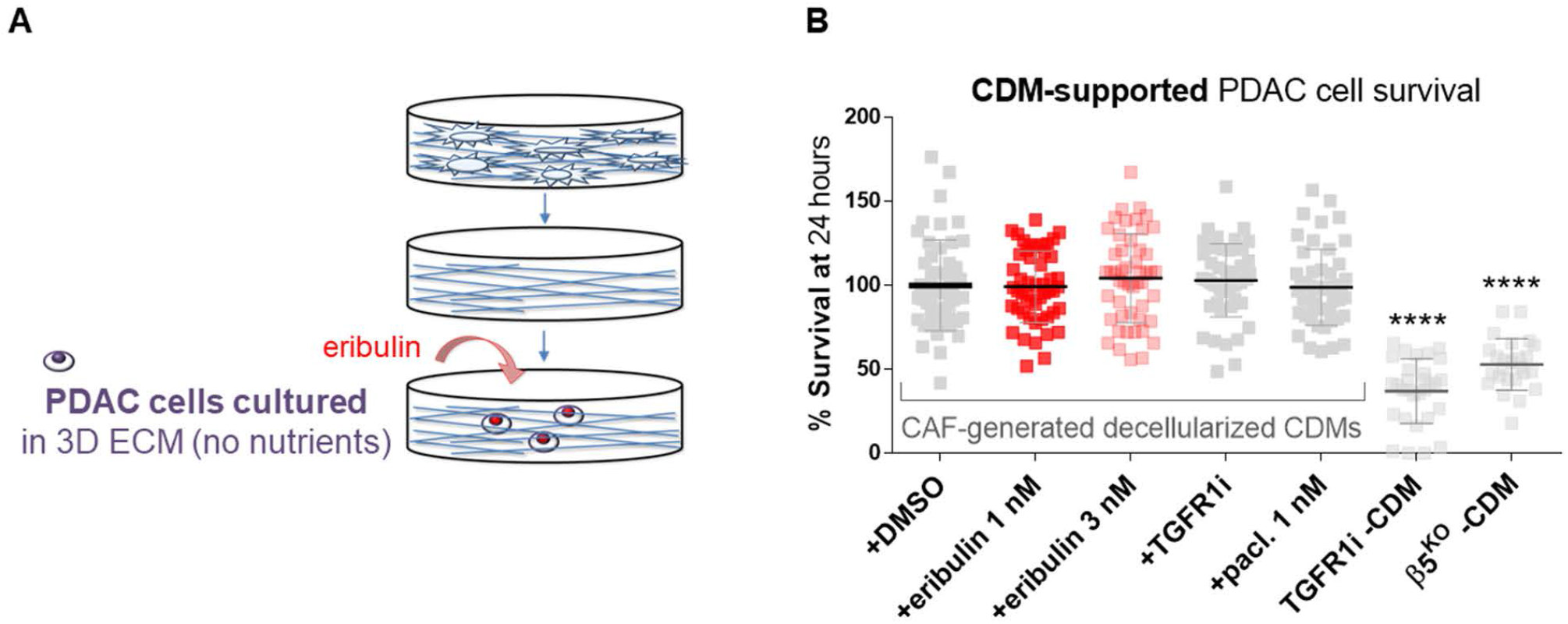
Eribulin fails to reverse the survival benefit provided by CAF ECMs. (A) Illustration showing the used experimental conditions whereby eribulin was added to nutrient-deprived cancer cells cultured within intact CAF-generated decellularized CDMs. (B) The graph shows the survival percentage of RFP-expressing Panc-1 cells treated with DMSO, eribulin, TGFR1i, or paclitaxel cultured for 24 hours in intact CAF-generated CDMs. CDMs obtained from TGFR1i treated and β5^KO^ CAFs (as in Figure 4C) served as “normalized” CDM controls. Data were normalized to DMSO veh., which average was set as 100% survival. For statistical significance, a one-way ANOVA (Turkey’s multiple comparisons) test was used to compare all conditions. Black asterisks denote comparisons to DMSO, while the eribulin vs paclitaxel comparison is noted in orange. * P < 0.05; ** p < 0.01; *** P <0.001; **** p < 0.0001.

## Discussion

In addition to being used as a third-line treatment option for patients with metastatic breast cancer [42], it was reported that eribulin prompts an overall survival benefit in a sarcoma Phase III clinical trial [43]. Importantly, several advanced PDAC patients treated with eribulin presented with a significant period (∼nine months) of stable disease [44]. This achievement is remarkable for this patient population, and therefore it justified the further investigation of using eribulin in this disease [18, 44]. These types of studies displayed encouraging efficacy for eribulin treatment in cancers that are fibrous (e.g., sarcomas) or that typically encompass a dense fibrous desmoplastic TME (e.g., advanced breast and PDAC). The above-stated facts highlight that in addition to its known microtubule-fixing function known for targeting epithelial cancer cells, eribulin also effectively regulates fibrous tumors like sarcoma. Further, eribulin mesylate was reported to be both a non-taxane microtubule dynamics blocker [38], as well as an EMT inhibitor that reinstitutes epithelial characteristics to mesenchymal cells via limiting Smad activity [20]. Of note, Smad activation lies downstream to TGFR1 canonical signaling, induced by the activated/mature TGFβ ligand [45]. It is hence not surprising that eribulin was shown to rapidly inhibit TGFβ-induced signaling [39]. Consequently, in this study, we hypothesized that eribulin could affect fibroblastic cells, which also greatly depend on TGFβ signaling for their activation [46]. The goal of this study was hence to assess the *in vitro* modulating effects that eribulin imparts on PDAC CAF function. For this, we used our well-characterized *in vivo*-mimetic desmoplastic system, referred to in this study as the CAF-generated CDM system, which is ideally suited for testing CAF phenotypes and functions [15, 23, 40].

By incorporating human CAFs with CAF-generated ECM, the above-mentioned CDM system was developed to simulate human desmoplasia [28, 34, 40]. The CDM is a pathophysiologic-relevant system that has been successfully used in numerous human cohorts [15, 23, 47]. The desmoplastic/CAF CDM system serves for studying drug effects on cancer regulated by the TME [30, 31, 48], as well as for measuring the direct influences of drugs on modulating fibroblastic cell function [22, 23, 29, 33]. Since quiescent/normal fibroblasts are naturally tumor-suppressive [10] and because CAF elimination has been demonstrated to be advantageous to PDAC progression and deleterious to patients [7-9], the field has placed significant efforts on identifying potential stroma-normalizing drugs [6, 12, 13].

To study the potential of drugs to impart a normalizing function upon CAFs, an important control to incorporate is a bona fide “normalized” CAF cell line. To this end, β5^KO^ CAFs constitute an ideal human cellular model as these cells present with a phenotype and function akin to perpetuating TGFβ inhibition [15]. In fact, β5^KO^ CAFs have been reported to generate CDMs with tumor-suppressive functions and traits [15, 22, 33]. Using β5^KO^ CAFs, we previously demonstrated that canonical TGFβ signaling is necessary for the production of tumor-supportive ECMs. However, once these matrices are produced, normal fibroblast undergo tumor-suppressive to tumor-supportive activation in response to these pro-tumoral natural scaffolds/substrates in a TGFβ independent manner [15]. The main difference between pharmacologically TGFβ-inhibited CAFs and β5^KO^ CAFs is that the latter perpetuates a tumor-suppressive “normalized” CAF phenotype and function, while the former is transient. This noted difference could explain the reason that results from experiments conducted in this study using eribulin, phenocopied β5^KO^ CAFs better than TGFR1i-treated CAFs.

While it is well accepted that TGFβ inhibition effectively blocks pro-tumor CAF activation, anti-tumor immunity, and EMT [15, 24, 26, 49], drugs targeting TGFβ have not been successful in clinical trials [16]. Considering that the level of eribulin toxicities is well tolerated in patients [18, 50], and knowing that eribulin limits canonical TGFβ signaling [20, 39], we posited that eribulin acts as a stroma-normalizing molecule. Results obtained from this study showed that eribulin causes increased CAF ECM fiber isotropy, as well as decreased CAF palladin, pFAK, pSmad2/3, and activated α5β1-integrin. These traits are all indicative of a tumor-supportive to tumor-suppressive CAF transition [15, 22, 27, 40]. Moreover, CAF-generated CDMs produced in the presence of eribulin lose their ability to induce CAF activation (e.g., failed to promote the reported αSMA localization to stress fibers [15, 27]). Importantly, while CAF-generated CDMs imparted modest protection against eribulin upon PDAC cells, eribulin treatment effectively “normalized” CAF-generate CDMs, which fail to support invasive spread as well as to effectively nurture PDAC cells under nutrient-deprived conditions [22, 23, 33, 35]. These results shed some light in potentially explaining the positive results in using eribulin specifically on fibrous-relevant cancers. Further, since eribulin effectively simulates TGFβ downregulation [24, 25], which has been shown to reinstitute anti-tumor immunity [26], results from this study also support the idea of combining eribulin with immune-regulatory drugs in the future.

In conclusion, data generated in this study suggest that eribulin simulates TGFβ inhibition by rendering CAFs with tumor-suppressive traits and functions (Figure 6).

**Figure 6.**
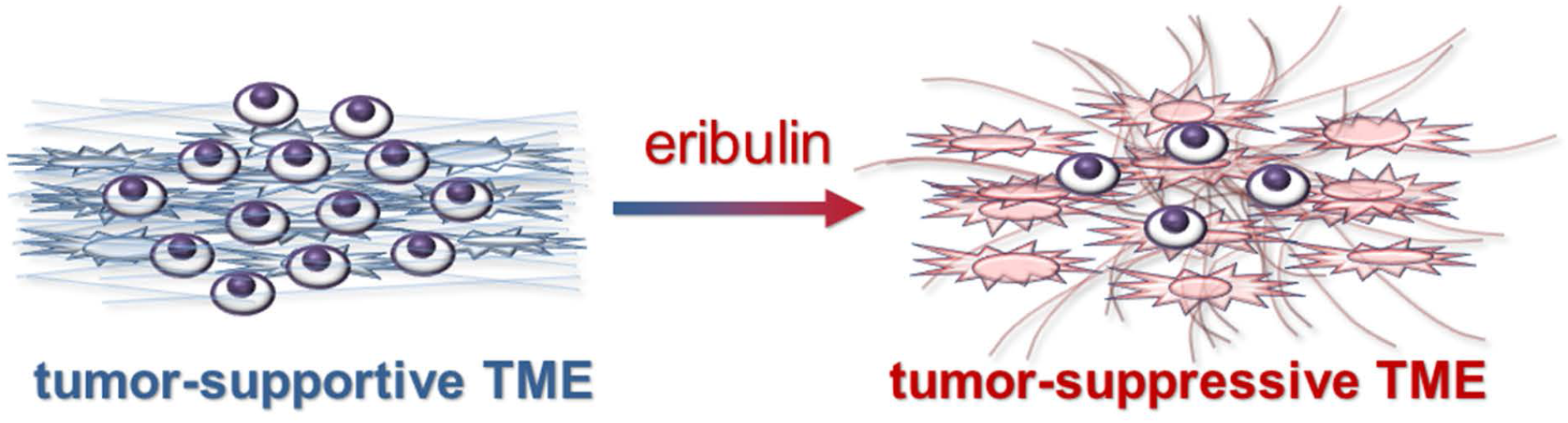
Schematic representation illustrating the tumor-suppressive effects of eribulin through normalization of CAFs. Illustration showing a tumor-permissive microenvironment (blue) and indicating that eribulin modulates CAFs in a way that these cells are prompted to generate a normalized, tumor-suppressive(red), TME.

## Acknowledgments

The authors like to dedicate this study to the memories of Dr. Patricia Keely, who revolutionized the desmoplastic ECM field, and Neelima Shah, who remains an inspirational figure to our team. We thank Drs. Janusz Franco-Barraza, Débora B. Vendramini-Costa, Ralph Francescone, Kristopher Raghavan, Jaye Gardiner, and Charline Ogier for their noteworthy comments throughout the experimental and manuscript preparation periods. This study was supported by funds obtained from Eisai Inc., as well as by the Pancreatic Cancer Cure Foundation, Pennsylvania’s DOH Research Formula Funds, the 5th AHEPA Cancer Research Foundation, Inc., a grant by the Worldwide Cancer Research, the Concetta Greenberg Pancreatic Cancer Institute Funds, and NIH/NCI grants R21CA231252, R21CA252535, R01CA232256, and the Core Comprehensive Cancer Center Grant P30CA06927 in support to Fox Chase Cancer Center’s facilities including BioSample Repository, Histopathology, Light Microscopy, Biostatistics and Bioinformatics, Immune Monitoring, and Cell Culture.

## Availability of data and materials

The datasets used and/or analyzed during the current study are available from the corresponding author on reasonable request.

## Authors’ contributions

Both authors were involved in the preparation of this manuscript. EC designed the experiments, wrote and supervised the writing of the manuscript. TL designed the experiments, carried out experiments, collected the data, and wrote the manuscript. Both authors read and approved the final manuscript.

## Ethics approval and consent to participate

All fibroblastic human cells used in this study were in accordance with guidelines and regulations under the approval of Fox Chase Cancer Center’s Institutional Review Board. To assure the protection of patient identities, samples were decoded before distributed to the research team. See Materials and Methods for more detail.

## Consent for publications

Not applicable.

## Supplemental Legends

**Supplemental Figure 1. Eribulin normalizes CAF traits**. (A) cellularity (left), measured as nuclei area coverage per image, and ECM thickness (right) are shown for the assorted conditions. (B) Graphs obtained from matrices generated by three independent CAFs, cultured as in Figure 2, indicating levels of fiber alignment, quantified as the percentage of fibers oriented within 15° from the mode angle. (C) Quantifications obtained from immunoblots of lysates collected at the end of matrix production, as in Figure 2, using an additional human pancreatic CAF, indicating levels of palladin and pSmad2/3 (GAPDH was again used as intracellular protein loading control). For statistical significance, one-way ANOVA was used to compare to DMSO vehicle. * P < 0.05; ** p < 0.01; *** P <0.001; **** p < 0.0001.

**Supplemental Figure 2. Eribulin-treated CAFs generate ECMs that fail to sustain PDAC cell survival under nutritional stress**. RFP-expressing Panc-1 cells from Figure 4C were cultured into the assorted ECMs using nutrient-depleted conditions. Viable cells were gauged following 48, and 72 hours as area coverage. Data were normalized to DMSO veh., which average was set as 100% survival. For statistical significance, a one-way ANOVA (Turkey’s multiple comparisons) test was used to compare all conditions. Black asterisks denote comparisons to DMSO, while the eribulin vs paclitaxel comparison is noted in orange. * P < 0.05; ** p < 0.01; *** P <0.001; **** p < 0.0001.

