## supplemental figures for "Eribulin normalizes pancreatic cancer-associated fibroblasts by simulating TGFβ inhibition"

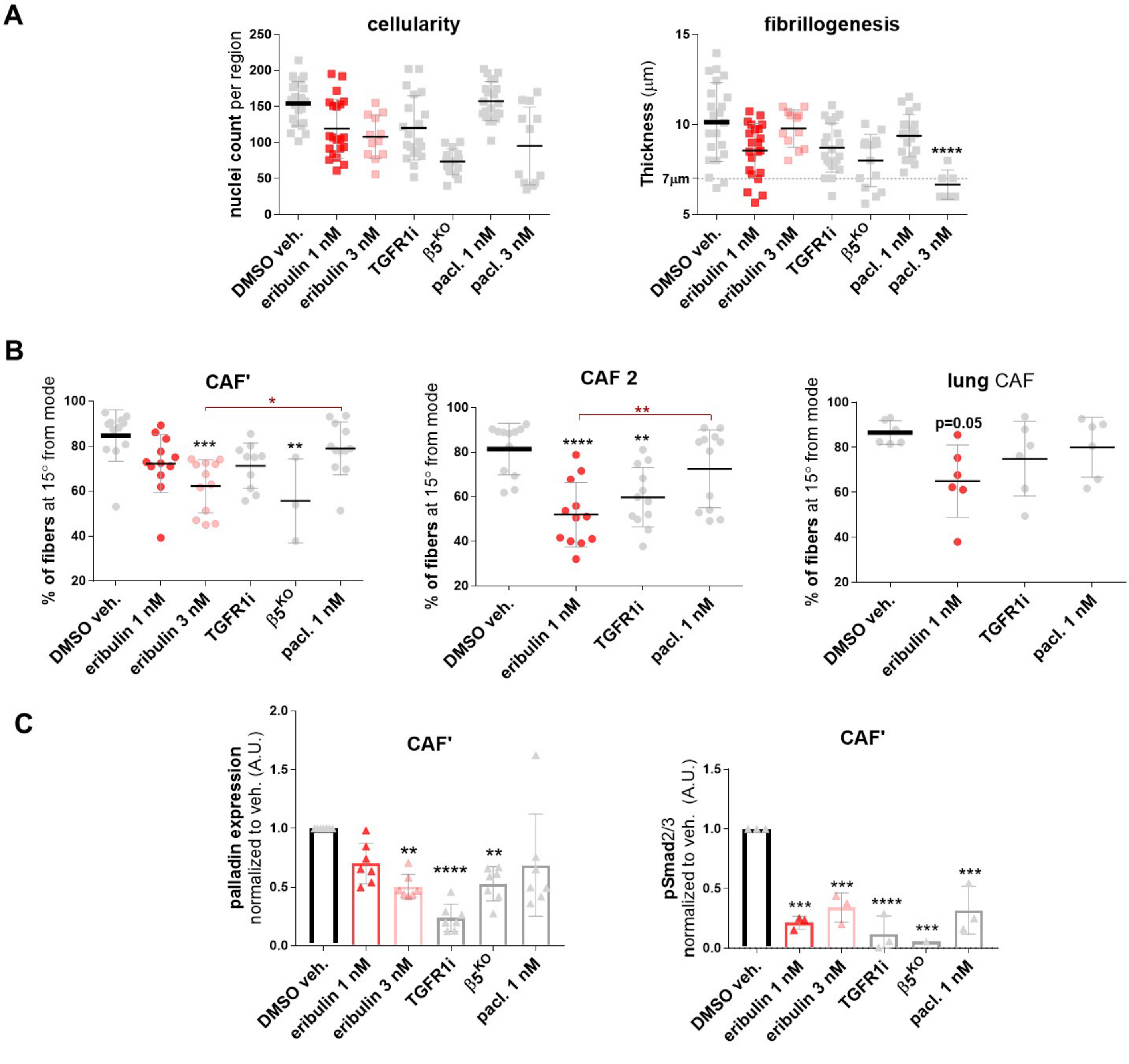


**Supplemental Figure 1. Eribulin normalizes CAF traits**. (A) cellularity (left), measured as nuclei area coverage per image, and ECM thickness (right) are shown for the assorted conditions. (B) Graphs obtained from matrices generated by three independent CAFs, cultured as in Figure 2, indicating levels of fiber alignment, quantified as the percentage of fibers oriented within 15° from the mode angle. (C) Quantifications obtained from immunoblots of lysates collected at the end of matrix production, as in Figure 2, using an additional human pancreatic CAF, indicating levels of palladin and pSmad2/3 (GAPDH was again used as intracellular protein loading control). For statistical significance, one-way ANOVA was used to compare to DMSO vehicle. * $P < 0.05$; ** $p < 0.01$; *** $P < 0.001$; **** $p < 0.0001$.

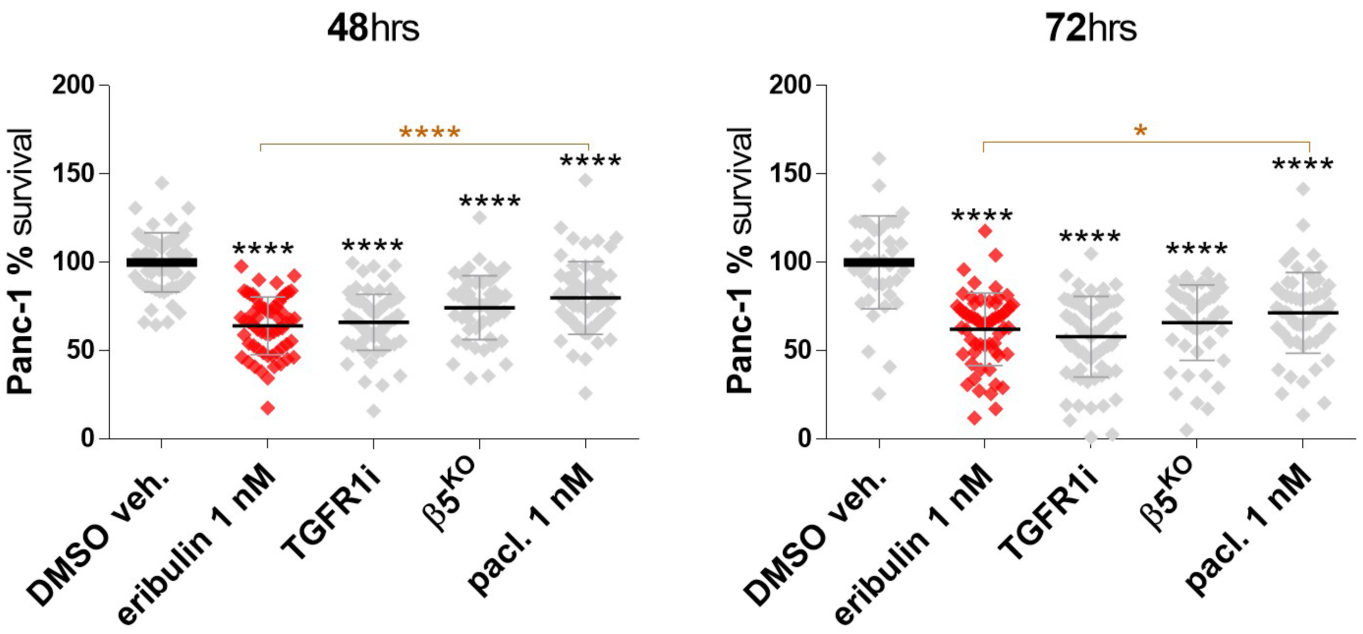


**Supplemental Figure 2. Eribulin-treated CAFs generate ECMs fail to sustain PDAC cell survival under nutritional stress.** RFP-expressing Panc-1 cells from Figure 4C were cultured into the assorted ECMs using nutrient-depleted conditions. Viable cells were gauged following 48, and 72 hours as area coverage. Data were normalized to DMSO veh., which average was set as 100% survival. For statistical significance, a one-way ANOVA (Turkey's multiple comparisons) test was used to compare all conditions. Black asterisks denote comparisons to DMSO, while the eribulin vs paclitaxel comparison is noted in orange. * $P < 0.05$; ** $p < 0.01$; *** $P < 0.001$; **** $p < 0.0001$.
